# Powerful decomposition of complex traits in a diploid model using Phased Outbred Lines

**DOI:** 10.1101/042176

**Authors:** Johan Hallin, Kaspar Märtens, Alexander I. Young, Martin Zackrisson, Francisco Salinas, Leopold Parts, Jonas Warringer, Gianni Liti

**Author notes:** Correspondence to: Gianni Liti or Jonas Warringer or Leopold Parts. Current address: Millennium Nucleus for Fungal Integrative and Synthetic Biology (MN-FISB); Departamento de Genética Molecular y Microbiología, Pontificia Universidad Católica de Chile, Casilla 114-D, Santiago, Chile. These authors contributed equally to this work.

## Abstract

Explaining trait differences between individuals is a core but challenging aim of life sciences. Here, we introduce a powerful framework for complete decomposition of trait variation into its underlying genetic causes in diploid model organisms. We intercross two natural genomes over many sexual generations, sequence and systematically pair the recombinant gametes into a large array of diploid hybrids with fully assembled and phased genomes, termed Phased Outbred Lines (POLs). We demonstrate the capacity of the framework by partitioning fitness traits of 7310 yeast POLs across many environments, achieving near complete trait heritability (mean *H*^2^ = 91%) and precisely estimating additive (74%), dominance (8%), second (9%) and third (1.8%) order epistasis components. We found nonadditive quantitative trait loci (QTLs) to outnumber (3:1) but to be weaker than additive loci; dominant contributions to heterosis to outnumber overdominant (3:1); and pleiotropy to be the rule rather than the exception. The POL approach presented here offers the most complete decomposition of diploid traits to date and can be adapted to most model organisms.

---

Decomposing the trait variation within natural populations into its genetic components is a fundamental goal of life sciences but has proven challenging^1,2^. Environmental and gene-by-environment influences are difficult to control for and alleles accounting for trait variation tend to have frequencies that are too low for their mostly weak effects to be reliably called^3^. Compounding matters, many alleles are believed to influence each other within (dominance) and between (epistasis) loci^4^. Consequently, traits depend on the presence of very many allele combinations that are exceedingly rare and whose contributions are near impossible to assess^5^. Model organisms offer more complete dissection of complex traits because they can be analyzed in controlled contexts, minimizing environmental variation, and in populations derived from a few founders, ensuring high frequencies of all alleles and allele combinations^6,7^. Because of their ease of use in genomics^8^ and phenomic screening^9^ large panels of haploid yeast segregants have allowed for exhaustive dissection of complex traits^10-12^. Unfortunately, exhaustive trait decomposition in haploid crosses requires the costly genotyping of thousands of genomes, disregards dominance, and provides much simplified estimates of epistasis. A more complete partitioning of trait variation that is relevant to a diploid context has therefore remained elusive. We here introduce a powerful and cost-effective framework for tracking the covariation through genome and phenome that allows accurate estimates of dominance and epistasis in diploid models. The framework is based on intercrossing two natural genomes over many sexual generations to reduce linkage^13,14^ followed by sequencing and systematic pairing of the resulting haploid recombinant segregants to generate a very large array of diploid hybrids with fully assembled and phased genomes, termed Phased Outbred Lines (POLs). We validate the capacity of the POLs approach by genetic decomposition of trait variation across 7310 diploid yeast genomes.

## A novel experimental framework for exhaustive analysis of diploid traits

To accurately decompose diploid traits, we isolated and sequenced the full genomes of 12^th^ generation haploid offspring. These haploid offspring were randomly drawn from a highly intercrossed two-parents founder pool with short linkage regions (**Fig. 1a**). The founder pool was derived from two highly diverged (0.53% nucleotide difference) wild yeast strains, here termed North American (NA) and West African (WA), such that only two alleles segregate with equal representation in the pool on average^13^. 172 haploid offspring from this pool, representing both mating types equally, were systematically crossed in all possible pairwise combinations to generate 7310 genetically unique diploid hybrids (POLs, **Fig. 1a**, Online Methods). With only a modest number of 172 haploid genomes sequenced^15^, we could accurately infer the genomes of our large set of 7310 POLs. Importantly, these genomes are fully phased, i.e. we know which parent contributed with which alleles. Furthermore, they had a very small fraction of missing genotype information (max: 6.5%; mean: 0.5%; median: 0.1%; min 0%) and there are no confounding effects from segregating auxotrophies that contribute to trait variation (**Supplementary Data SI**). The hybrids showed remarkable uniformity, with heterozygote frequencies close to 50% (**Fig. 1b**). The few strong deviations (8 deviations > 30%) from 50% heterozygosity were either due to selection for one parental allele during the intercross (overrepresentation of homozygous sites) or from the crossing design, the latter resulting in regions of fixed heterozygosity at the *MAT* and *LYS2* loci (**Fig. 1b**).

**Figure 1.**
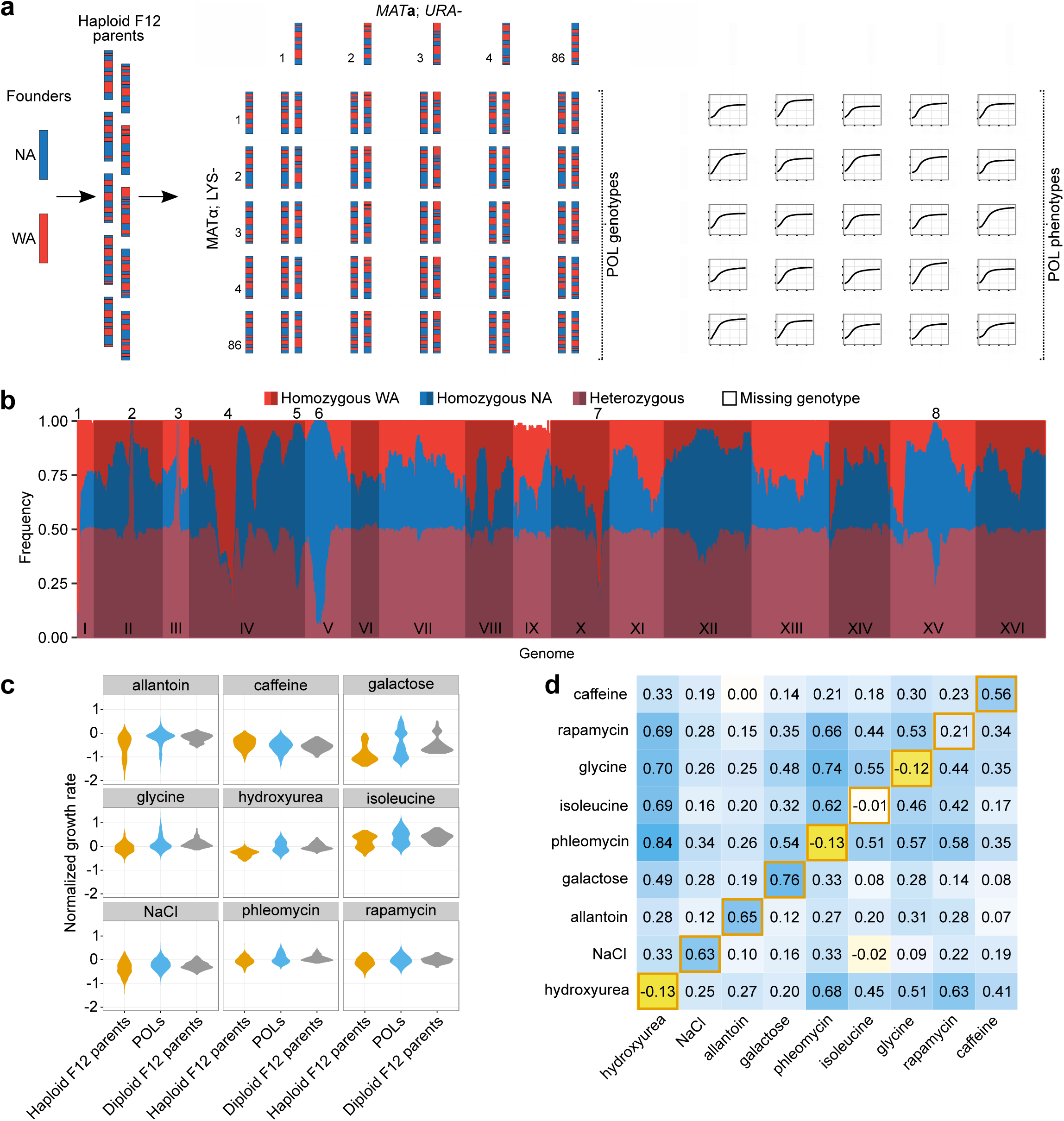
A novel experimental framework for high-powered analysis of diploid traits. (**a**) Experimental design. Left panel: Advanced intercrossed lines were construceted by multiple rounds of random mating and sporulation of North American (NA) and West African (WA) genomes. Midlle panel: We sequenced 172 of the resulting segregants and paired these to generate an array of 7310 diploid hybrids (POLs). Right panel: The POLs and their F12 haploid parents were growth phenotyped in nine environments, providing high resolution growth curves. (**b**) Frequency of homozygotes (red: WA/WA, blue: NA/NA), heterozygotes (purple: NA/WA) and missing genotypes (white, mostly attributed to chr. IX aneuploidies) at each segregating site among the 7310 POLs. Deviations from 50% heterozygosity are explained by selection (numbers 1, 4-8) against one allele in the F12 haploid parent construction, or by forced heterozygosity at the *LYS2* (number 2) and *MAT* (number 3) loci. (**c**) Maximum growth rate distributions of POLs (blue), their haploid F12 parents (orange) and the diploid parent estimates (gray, Online Methods). (*d*) Correlations (Pearson’s *r*) between the maximumgrowth rate and total growth for POLs within environments (lower left to upper right diagonal; orange boarders),between maximum gorwth rates (above diagonal) and total growth (below diagonal) in pairs of environments. Color intensity (3-color scale: dark yellow to white to dark blue) and number indicate the degree of correlation (r)

We precisely phenotyped the complete set of 7310 designed POLs (median CoV = 10%, mean CoV = 14%), their F12 haploid parents, the diploid NA and WA founders and their hybrid in a well replicated (*n*≥4), high resolution growth phenotyping platform designed to minimize noise and bias^16^. We selected nine physiologically distinct environmental conditions (**Supplementary Table SI**) that challenged growth to different extents (**Supplementary Fig. S1a**), and we obtained more than 56 million population size estimates, organized into 250.000 growth curves (**Fig. 1a**, **right panel**). Extracting the maximum growth rate (population doubling time) and total growth (total change in population size) from each growth curve (**Supplementary Data SII**), we found phenotype distributions across the POLs to be mostly monomodal (**Fig. 1c** **and Supplementary Fig. S1b**). Given the absence of environmental variation, this implies complex traits with multiallelic influences. Galactose and allantoin growth were bimodally distributed, in agreement with large effect sizes for the *GAL3* (WA premature stop codon) and *DAL* (linked loci, WA loss-of-function SNPs in *DAL1* and *DAL4*) genes respectively^17,18^. Correlation between maximum growth rate and total growth ranged from −0.13 to 0.76 (Pearson’s *r*; **Fig. 1d, orange borders**) but was overall low (mean *r*: 0.27; median *r*: 0.21), in agreement with the hypothesis that distinct genetic factors control population expansion in different growth phases^17,19^. Correlations across environments were positive in all but one case (*r*=-0.02) and often of moderate or large magnitude (max *r*=0.84, median *r*=0.29; **Fig. 1d**). We cannot completely exclude a small influence of shared error on correlations, but the extensive standardization, randomization and normalization (Online Methods), and the large variation in pairwise correlations argue compellingly in favor of extensive positive pleiotropy.

## Near complete decomposition of diploid traits into their genetic components

Based on the assembled and phased diploid genomes, we used a random effects model to decompose the variance in growth traits into components arising from additive (no interaction), dominance (intralocus interaction) and pairwise and third order epistatic effects (interlocus interactions) (**Supplementary Note SI**). The large sample size and known large variation in relatedness allowed us to estimate nonadditive variance components with unprecedented accuracy (mean S.E.M = 1.0-2.5%, depending on environment). Simulations provided standard errors of the mean in the same range (**Supplementary Table SII and Supplementary Data SIII**). The large number of replicates and the accuracy of the normalized phenotype measures minimized environmental variation. Therefore, additivity, dominance, and pairwise epistasis accounted for almost all trait variation ([80-99% depending on environment], median 90%, **Fig. 2, upper panel**). On average, the proportion of phenotypic variance explained by additive effects was 74% (50-87%), for dominance effects this was 9% (2-31%), and for pairwise interactions this was 8% (1-15%). We estimated that third order interactions accounted for 1.8% of the trait variation on average (**Fig. 2, lower panel**), but only maximum growth rates on isoleucine and glycine and total growth in presence of phleomycin were significantly (>2 S.E.M from 0) affected by third order epistasis. Complete dominance of the functional NA over the nonfunctional WA cluster of *DAL* genes^18^ ensured a large dominance component for variation in allantoin growth rate and total growth. Otherwise, the large variance contributions of additive genetic influences were consistent across environments (**Fig. 2, upper panel**). The trait with the largest estimated variance from pairwise epistasis was growth rate on glycine (15%); this epistasis variance contribution equaled half of the largest dominance variance estimate (31% for allantoin growth rates). Variation in genome wide levels of homozygosity had no detectable influence on yeast fitness traits (**Supplementary Fig. S2**). This is in stark contrast to its substantial negative effect on human traits, e.g. height^20^). Thus, the data suggest that there is no general inbreeding depression in yeast, consistent with natural populations being largely homozygous^21,22^.

**Figure 2.**
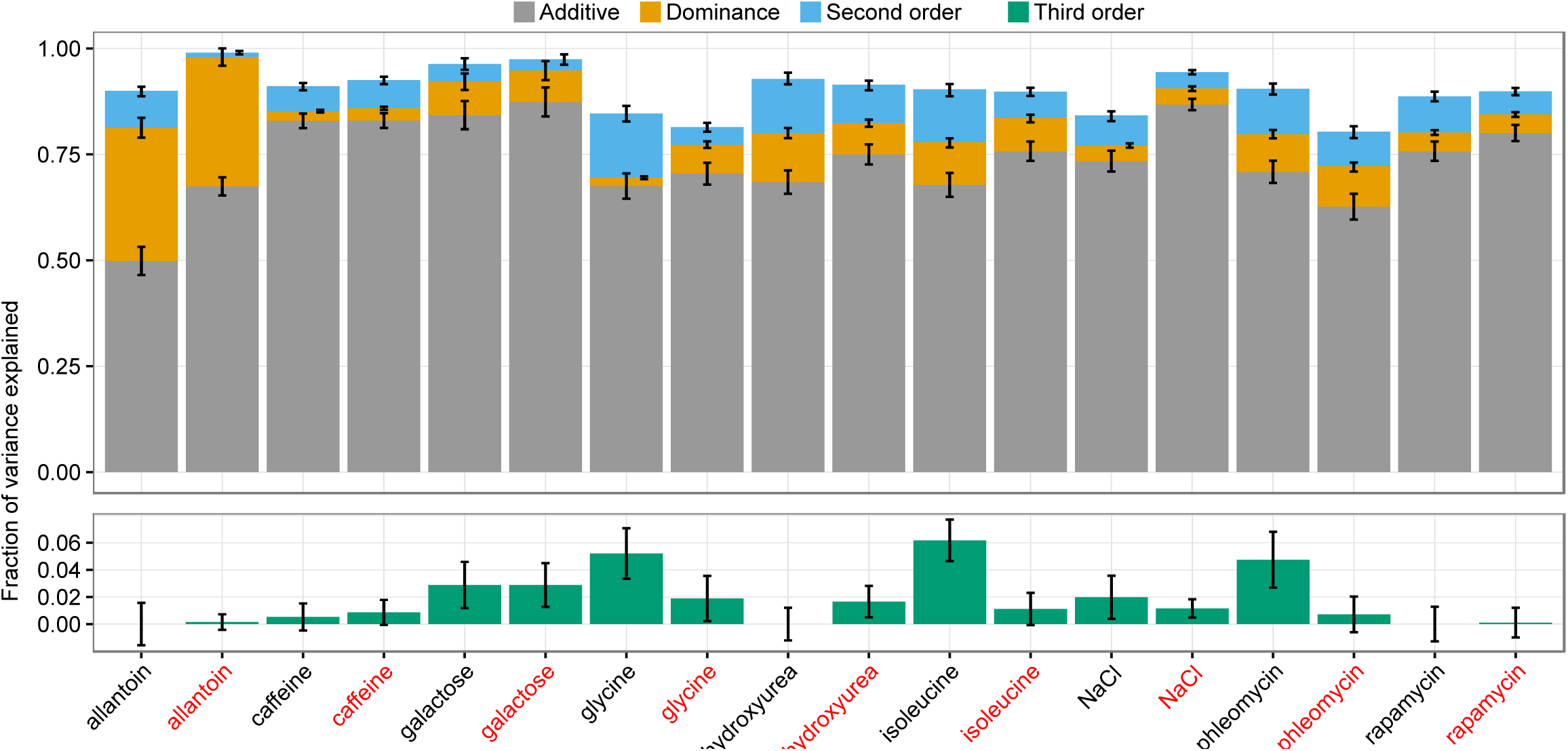
Near complete decomposition of diploid traits into their genetic componenets. DEcomposing the total variance in growth traits across 6642 diploids into additive (gray, upper panel), dominance (yellow, upper panel), second order epistatic (blue, upper panel) and third order epostatic (green, lower panel) genetic contributions. Black label = growth rate, red label = total growth. Error bars = S.E.M

## Cost-efficient QTL mapping in yeast POL diploid hybrids

Our crossing design results in half the genome of each POL being kept constant across the 86 POLs that are derived from any one haploid F12 parent (**Fig. 1a**). This sharing of half a genome accounted for surprisingly much of the overall variation in traits^23^, which somewhat restricted our capacity to distinguish contributions from individual alleles and allele pairs from the effect of the genetic background. Nevertheless, our platform provided a cost-efficient framework for calling both additive and nonadditive (dominance and epistasis) QTLs in diploid models. We excluded 668 POLs with chr. IX aneuploidies and then mapped QTLs onto the inferred parent phenotypes (additive effect of genetic background) and recorded deviations from inferred parent means (nonadditive effects; Online Methods). Both QTL mapping approaches accounted for the population structure. We called a total of 145 unique QTLs at 10% false discovery rate (FDR). These included the *GAL3* stop codon, as well as the *DAL1* and *DAL4* non-synonymous and stop codon mutations, known to account for most of the variation in galactose and allantoin growth respectively (**Fig. 3a, Supplementary Fig. S3 and Supplementary Table SIII**). Some (21%) of the QTLs contributed significantly to both additive and nonadditive phenotype components, but the majority were private to one of them (**Fig. 3b**). The nonadditive (75%) outnumbered the additive (25%) QTLs, but explained on average less of the trait variation (6% vs. 28%, Student’s t-test: *p* = 2e-6, **Fig. 3c**). Thus, significant nonadditive trait contributions were more common but weaker. The QTLs were confirmed using linear mixed models that separated additive, dominant and epistatic effects (Online Methods). In almost all cases, nonadditive QTLs coincided with dominance effects (**Fig. 3a**). The complete recessiveness of the WA *GAL3* allele for galactose growth and of the WA *DAL* alleles for allantoin growth recapitulated established knowledge^17,18^ (**Supplementary Fig. S4a**).

**Figure 3.**
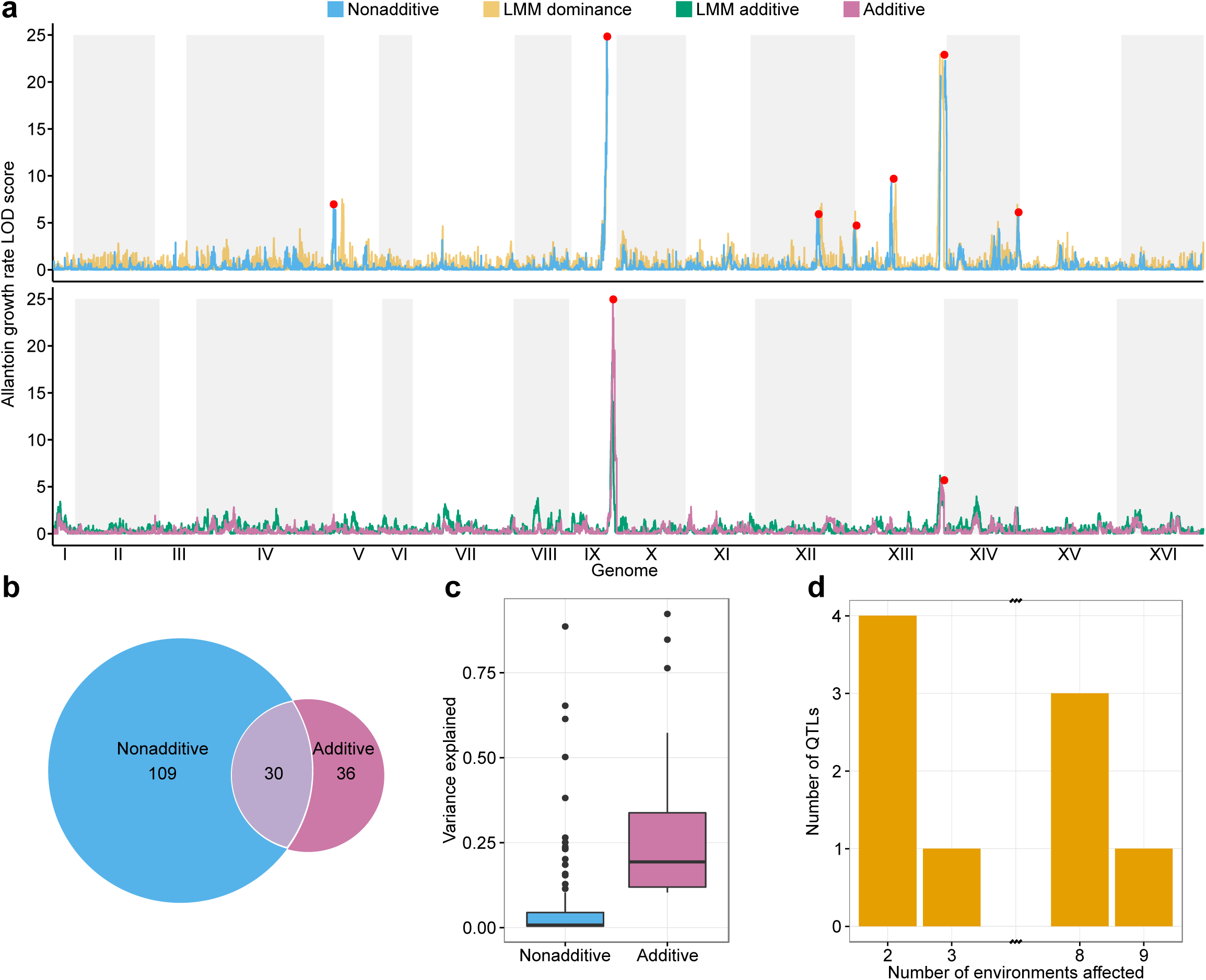
Cost-efficient QTL mapping in yeast POLs. QTLs were mapped across 6642 genomes and 18 traits based on additive and nonadditive contributions. QTLs were validated as additive or dominant genetic contributions using Linear MIxed Models (LMM). (a) QTL signal strength (LOD score, *y*-axis) as a function of genomic position (*x*-axis), for maximum growth rate on allantoin as sole nitrogen source, using additive (LMM and non-LMM; lower panel) and nonadditive (non-LMM and LMM only capturing dominance; upper panel) models. Red dots indicate significant (FDR, *q* = 10%) QTL calls. White/grey fields indicate chromosome spans. (**b**) Venn diagram of significant QTLs capturing additive and nonadditive genetic contributions. All 18 (growth rate and total growth over nine environments) were considered, with pleiotropic QTLs counted multiple times. (**c**) Tukey boxplot showing the fraction of total variance in a trait explained by additive (purple) and nonadditive (blue) significant QTLs (non-LMM models). (**d**) Histogram of pleiotropic QTLs. A QTL was counted as shared across environments if peaks were within 10 kb of each other. No QTLs were significant in 4, 5, 6 or 7 environments.

Only 32 of 145 (22%) additive and nonadditive QTLs called were unique to a single environment, reflecting that extensive pleiotropy is the rule rather than the exception (**Fig. 3d**). Almost half (44%) of the pleiotropic QTLs affected at least five environments, with universal growth QTLs on chr. XIII penetrating regardless of the environment and one QTL on each of chr. IX, X and XV penetrating in all but one environment (**Fig. 3d**). Given the wide span of environmental effects on growth and cellular physiology in our set of environments, this prevalence and penetrance of universal growth QTLs is remarkable. More QTLs (69%) were shared between maximum doubling time and total growth than expected from their low general correlation (mean *r*: 0.27, **Fig. 1d**). This was largely explained by the near universal chr. IX QTL affecting the two fitness components antagonistically: NA homozygotes grew slower but reached higher total growth (**Supplementary Fig. S4b**). This profound fitness trade-off penetrated regardless of environment and may therefore have had a large influence on natural selection on the ancestral wild strains. Finally, we note that disproportionately many (28% vs. 9% expected, Fisher’s exact test, *p* < 0.0001) QTLs were subtelomeric; almost all (84%) of these were pleiotropic. This agrees with previous haploid studies, and adds credibility to the suggestion that hypervariable subtelomeres account for much of the remarkably large trait variation in yeast^24,25^.

## Explaining parent-offspring discordance by intralocus interactions

The degree to which offspring phenotypes deviate from the mean of the parent phenotypes, heterosis, and which genetic factors account for this difference are central questions in breeding. Capitalizing on the scale (130.000 offspring traits) of our screen, we established the phenotype discordance of the POLs from their diploid parents (Online Methods) with previously unattainable completeness. Of the offspring phenotypes that differed significantly from those of both its parents and allowed discordance analysis (**Supplementary Fig. S5a**), the majority (88% to 92%) deviated significantly from the midparent expectation and were thus heterotic. Half (41 to 52%) of these cases corresponded to the offspring being either superior (best parent heterosis) or inferior (worst parent heterosis) to both parents, with equal prevalence of best parent and worst parent heterosis (**Fig. 4a**). This is surprising given that smaller scale studies have indicated higher prevalence of best parent heterosis^26,27^.

**Figure 4.**
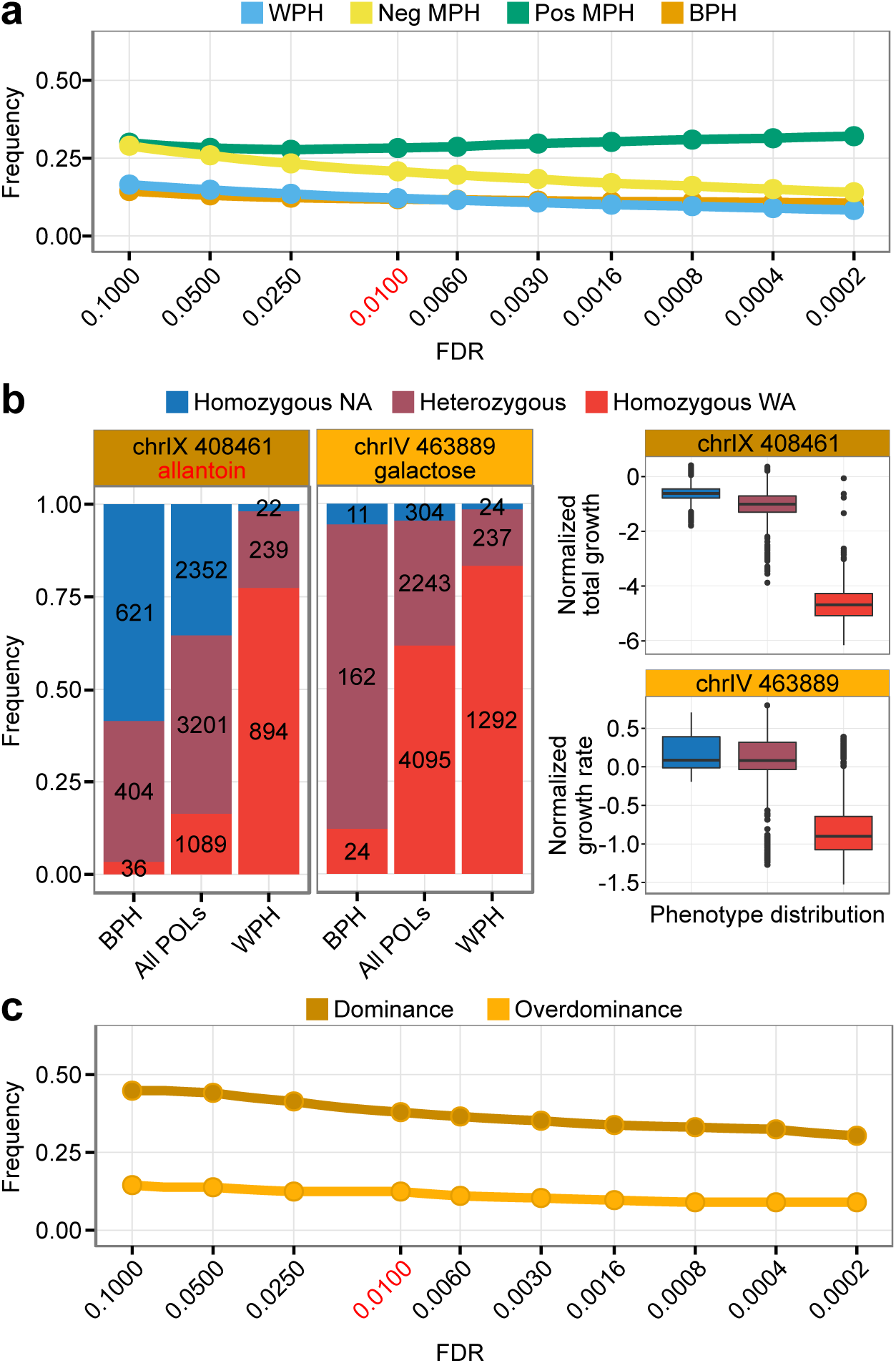
Explaining parent-offspring discordance by intralocus interactions. Abbreviations: worst parent heterosis (WPH); mid parent heterosis (MPH); best parent heterosis (BPH). (**a**) Frequencies of the heterotic POLS (*y*-axis) as a function of a range of FDR significance cut-off (*q*) values (*x*-axis). Line color = type of heterosis. Red text = FDR *q*-value chosen for downstream analysis (**b,c**). (**b**) Left panel: example of QTLs called as contributing to best parent heterosis by dominance (dark orange) and by overdominance (light orange) respectively. Dominance was called as enrichment of strongest homozygote and overdominance as enrichment of heterozygous state among BPH POLs as compared to all POLs (left panel). Right panel: phenotype (top: allantoin, bottom: galactose) distribution depending on genotype composition at the same QTLs. (**c**) The frequency of QTLs called as contributing by dominance and overdominance respectively (*y*-axis) as a function of FDR significance cutoff (*q*) values (*x*-axis). Red label = FDR *q*-value chosen for downstream analysis.

Overdominance (heterozygotes being superior to both homozygotes) and dominance (heterozygotes differing from the parent mean) can both contribute to best parent heterosis. However, calling such contributions is challenging: multiple effects often act in parallel and they rarely manifest in all genetic backgrounds. We conjectured that overdominance contributions to offspring superiority should manifest as enrichment of heterozygotes among POLs with best parent heterosis. Similarly, dominance contributions should manifest as enrichment of the better homozygote. For each QTL and trait separately, we therefore called overdominance contributions as more heterozygotes than expected among best parent heterotic POLs (*χ*^2^ test, *p* < 0.01; **Fig. 4b, dark orange**), and dominance contributions as more of the better homozygote than expected (**Fig. 4b, light orange**). Overall, for 38% of QTLs, we identified probable dominance contributions to best parent heterosis, and for 12%, we identified probable overdominance contributions. These proportions were consistent across a wide range of significance cutoffs (**Fig. 4c**). For the remaining 56% of QTLs, no significant contributions to the best parent heterosis were detected.

The dominance/overdominance contributions to the growth of the best parent heterotic POLs were often notably different from contributions in the entire population (**Fig 4b, bottom left vs. bottom right panels**). Indeed, of the 18 QTLs for which we detected overdominance in the best parent heterotic POLs, only two showed heterozygote phenotype averages being significantly superior to homozygote phenotype averages when the entire set of POLs was considered (Student’s t-test, *p*<0.01). This suggests that dominance-by-dominance or dominance-by-additive interactions potentiate the best parent heterosis, shifting dominant or additive loci to overdominant to create best parent heterosis in a minority of backgrounds. For the chr. IX QTL with a near universal fitness trade-off, heterozygotes were consistently enriched among offspring with superior growth rate but underrepresented among those with superior total growth. Conversely, homozygotes with the North American allele were underrepresented among offspring with superior growth rate, but overrepresented among those with superior total growth (**Supplementary Fig. S5b**). Thus, overdominance and dominance at this universal QTL both contributed to best parent heterosis for growth, but for different growth components. Finally, we called underdominant contributions to worst parent heterosis as more heterozygotes than expected among worst parent heterotic POLs. Overall, we found 12% of QTLs to contribute underdominantly to worst parent heterosis (**Supplementary Fig. S5c**). To our knowledge, this is the most exhaustive dissection of heterosis to date.

## DISCUSSION

Traits have been exhaustively mapped and decomposed in haploid models^10-12,28,29^ but extrapolation from haploid screens to the biology of diploids is precarious. Intralocus interactions in the form of dominance cannot be estimated in haploid screens and these also only capture additive-by-additive epistasis. Moreover, ploidy as such has a fundamental impact on traits^30^, both due to its influence on cell size and the masking of recessive alleles in diploids^31,32^. The Phased Outbred Lines (POLs) presented here circumvent the shortcomings of haploid screens by offering decomposition of diploid traits with previously unattainable exhaustiveness. The capacity of the approach follows from generating a very large array of fully phased diploid genomes based on short read sequencing of an only moderate number of haploids. The alternative, acquiring phased genomes from direct sequencing of diploids, would require long-read sequencing of thousands of isolates and will remain economically unfeasible even in model organisms for years to come^33^. As a direct consequence of the experimental design, each POL shares one haploid genome with siblings spawned from the same haploid parent. This sharing of half a genome had surprisingly large effects on trait similarity, greatly aiding both trait prediction from relatives^23^ and the partitioning of trait variation into its additive, dominant and epistatic components. In contrast, it somewhat restricted our ability to distinguish the weaker effects of individual loci and the calling of those QTLs. The large impact that sharing one haploid genome has on trait similarity among diploids, and the associated benefits and drawbacks, may or may not manifest in other model organisms. Beyond the removal of the sex-switch and introduction of sex-specific auxotrophic markers, POLs impose no requirements on the yeast genotypes used; the design is lineage agnostic. The diploid hybrids have identical marker composition, avoiding marker effects that confound many haploid crossing designs^34,35^. The framework allowed partitioning diploid trait variation into its major components with little room for confounding effects, due to nearly all trait variation being accounted for. Additive effects explained the vast majority of phenotypic variation, with approximately equal variance contributions from dominance and pairwise interactions at slightly less than 10%. The large explanatory power of additive genetics is well in line with findings in haploid screens^10,29^. Third order epistasis explained less than 2% of the trait variation, comparable to, or somewhat less than, estimated for third^11^, or third and higher^12^ order interactions in haploid yeast. Thus, although examples where three-way interactions affect trait variation can be found^12,36,37^, they generally account for little trait variation. Despite the lower overall contribution of nonadditive than additive genetics to trait variation, we found nonadditive QTLs to outnumber additive QTLs. The weaker mean effect of nonadditive QTLs partially explains this discrepancy. In addition, subtle differences in how QTLs were called means that we cannot completely exclude that we detected nonadditive effects with somewhat better power.

A stable haploid phase, indefinite storage as frozen stocks and easy mating will remain distinct advantages of yeast. Nevertheless, POLs can be employed in most higher model organisms, with only slight modifications to the approach. Panels of extensively recombined offspring can be generated using two or more founder parents in mouse, plants, flies and worm^38,39^. Successive inbreeding or selfing is common practice to produce recombinant inbred lines (RILs). The gametes of these sequenced RILs can be paired by designed mating to generate the final array of POLs to be phenotyped. To attain exhaustiveness while avoiding confounding effects from uncontrolled environmental variation, the cost-effectiveness of the genotyping needs to be matched by a phenotyping approach that achieves both scale and accuracy. The here reached broad sense heritability, with a lower bound mean estimate of 91%, may remain challenging to match in most species. Nevertheless, phenomics is advancing on broad fronts and simultaneous high throughput and accuracy is on the horizon in most model organisms^40^.

## METHODS

A full description of the methods is available in the online version of the paper.

## ACKNOWLEDGEMENTS

JH was supported by the Labex SIGNALIFE (ANR-11-LABX-0028-01) and KM was supported by the European Regional Development Fund through the BioMedIT project, FS was supported by ATIP-Avenir (CNRS/INSERM), Becas Chile, FONDECYT (3150156) and MN-FISB (NC120043) postdoctoral fellowships. This study was funded by the Swedish Research Council (325-2014-6547 and 621-2014-4605), the Research Council of Norway (222364/F20) to JW; by a Marie Curie International Outgoing Fellowship, the Wellcome Trust, and Estonian Research Council (IUT34-4) to LP; ATIP-Avenir (CNRS/INSERM), ARC (grant number PJA20151203273), FP7-PEOPLE-2012-CIG (grant number 322035), ANR (ANR-13-BSV6-0006-01 and Labex SIGNALIFE ANR-11-LABX-0028-01), Cancéropôle PACA (AAP émergence 2015) and DuPont Young Professor Award to GL.

## ONLINE METHODS

### Generation of Phased Outbred Lines

F12 outbred lines were derived from a two way intercross between ancestors of the North American (YPS128) and West African (DBVPG6044) populations, as described^13^. Ancestral strains differed at 0.53% of nucleotide sites^41^. Following random sporulation of F12 diploids, 86 haploids of each mating type were randomly selected and their mating type and auxotrophies determined. Haploid genotypes were selected to allow systematic crossing: *MAT***a**, *ura3∷KanMX*, *ho∷HygMX* and *MAT*α; *ura3∷KanMX*; *ho∷HygMX*; *lys2∷URA3*. Haploids of different mating types were robotically mated on rich medium (1% yeast extract, 2% peptone, 2% glucose, 2% agar) in all pairwise combinations combining their complementary *LYS* and *URA* auxotrophies using a RoToR HDA (Singer Ltd, UK). Diploid hybrids were selected twice on Synthetic Minimal (SM) medium (0.14% Yeast Nitrogen Base, 0.5% ammonium sulphate, 2% (w/v) glucose and pH buffered to 5.8 with 1% (w/v) succinic acid, 2% agar). The theoretical maximum amount of POLs from our experimental design was 7396 (86x86), however, one F12 haploid strain was contaminated prior to mating and all 86 hybrids spawning from this cross was therefore discarded (7310 were retained).

### Genotype construction

The haploid F12 parents were sequenced by short read sequencing^15^, and mapped to parental genomes in order to call segregating sites, infer genotypes and characterise the recombination landscape. All segregants were homoplasmic, carrying the non-recombined WA mtDNA genome. This excludes confounding mtDNA effects. Phased genomes of their diploid hybrid offspring was reconstructed *in silico* using custom R code. Diploid hybrids with chr. IX aneuploidies were excluded for the QTL mapping (6642 were retained).

### High resolution growth phenotyping

High resolution growth phenotyping on solid agar medium was performed using a 1536-colony plate layout. Each plate (Plus plate, Singer Ltd, UK) was cast with exactly 50mL of Syntetic Complete medium at 50C (as SM above with added 0.077% Complete Supplement Mixture (CSM, formedium)). Casting was performed on an absolutely leveled surface with drying for ~1 day. The base medium was supplemented with additional stressors or alternative carbon or nitrogen sources as indicated (**Supplementary Table SI**). The 7310 POLs were distributed over 1152 positions across eight plates. We used *n* = 4 replicates for each experimental plate, with replicates initiated from two different pre-cultures and run in different instruments and plate positions to minimize bias. Their 172 haploid F12 parents (*n* = 6 replicates on each plate, two plates) and their diploid NA and WA ancestral lineages (*n* = 72 replicates on each plate, two plates) were phenotyped separately. Every 4^th^ position was reserved for internal controls (diploid NA ancestral strains). These 384 controls were interleaved with experiments on pre-culture plates, ensuring equal treatment of controls and experiments. High resolution population size growth curves were obtained using Epson Perfection V700 PHOTO scanners (Epson corporation, UK) and the Scan-o-matic framework^16^. Scanners were maintained in a 30ºC, high humidity environment that minimized light influx and evaporation. Experiments were run for 72h, with automated transmissive scanning and signal calibration in 20 min intervals. Calibrated pixel intensities were transformed into population size measures by reference to cell counts obtained by flow cytometry. Raw population growth curves were slightly smoothed to remove noise. Poor quality curves (0.3%) were rejected following manual inspection. Retained population growth curves were broken down into two growth phenotypes i) maximum growth rate, extracted using linear regression from the steepest slope of the population’s exponential phase, and ii) total growth, extracted as the population size gain between the first (0h) and the last measure (72h). To capture spatial bias on each 1536 plate, the two growth phenotypes were normalized to the internal controls using the Scan-o-matic principle^16^. The final phenotypes used were the average phenotype across all replicates. Detailed protocols are available^16^. To circumvent the problem of calculating Coefficients of Variation (CoV) for normalized growth phenotypes spanning over both negative and positive values, these were reverted back into actual doubling times and yields, before CoV calculations. This reversion was performed by multiplying each normalized value with the median control trait value and reversion of the log transformation.

### Phenotype Variance Partitioning

We estimated additive relatedness from genotypes, derived formulae for covariance due to dominance, pairwise and third order interaction effects and fitted the model using restricted maximum likelihood, as in Yang *et al*. (2011)^42^. As third order interactions biased estimates of other variance components, these were analyzed and reported separately. Details are available in Supplementary Note SI.

### QTL mapping

QTL calling was made using the `scanone` function with the `marker regression` method in R/qtl^43^ with estimated diploid parent phenotypes (additive genetic background contribution to traits) and POL deviations from the estimated diploid parents values (variation not explained by additive effects of parental background) respectively using 52,466 markers. Diploid parental phenotypes were estimated as the median of all hybrids descended from that parent. Using the deviations from expected midparent phenotype for the POLs has the additional critical benefit of effectively accounting for population structure by removing the additive effect of the more similar genetic composition due to shared parents. Significance thresholds were given by permutations (x1000), 95% Bayesian credible intervals were calculated for each QTL using the `bayesint` function in R/qtl. QTL calling by linear mixed models, also accounting for population structure, was performed and used as verification. For these, in order to test each QTL, we constructed the realised genetic relationship matrix by discarding the SNPs within the 50kb neighbourhood of the SNP under consideration; these models were fitted with LIMIX^44^ as in Märtens, Hallin *et al*. (2015)^23^. Consecutive markers having the same genotype across all individuals were removed for increased computation speed, leaving 10,726 segregating sites^23^.

### Heterosis

We used a Student’s t-test to detect POLs significantly deviating (p < 0.01) from the mean parent phenotype, either overperforming (positive mid parent heterosis) our underperforming (negative mid parent heterosis). The parent phenotypes used were estimated from all POLs descending from the given parent as described under “QTL mapping” in Online Methods, the variance of the mean parent phenotype was set to equal that of the most variable parent. POLs deviating from the mean parent were then tested using a Student’s t-test (*p* < 0.01) for positive deviations from the strongest parent (best parent heterosis, BPH) and for negative deviations from the weakest parent (worst parent heterosis, WPH)

### Dominance, overdominance and underdominance contributions to heterosis

To test for overdominance contributions to best parent heterosis we compared the expected and observed number of heterozygous genotypes among best parent heterotic POLs (defined as above). This was performed for each QTL separately using a *χ*^2^ test. Entries were: observed number of heterozygotes and observed number of homozygotes (summed) among BPH POLs and the corresponding expected numbers, given distributions among all POLs. A range of cut-offs for significance was tested and the stability of results across cut-offs ascertained. We cannot completely exclude that pseudo-overdominance, i.e. tightly linked loci with dominance of opposite parental alleles, confuse some assignments of overdominance. However, given the small linkage regions, we expect pseudo-overdominance to be rare and the associated overestimation of overdominance to be small. We tested for dominance similarly, but pooling the weaker homozygote state with the heterozygote state and calling significant enrichment of the better homozygote among BPH POLs. Underdominance contributions to worst parent heterosis were called as for overdominance, but as enrichments of the heterozygous genotype among worst parent heterotic POLs.

## SUPPLEMENTARY INFORMATION

Supplementary Figure S1 | Growth phenotype distributions

Supplementary Figure S2 | Extent of genome-wide homozygosity has no impact on yeast growth

Supplementary Figure S3 | QTL maps for each trait

Supplementary Figure S4 | Phenotype distributions as a function of genotype for key QTLs

Supplementary Figure S5 | Dominance, overdominance and underdominance in heterotic POLs

Supplementary Table SI | Environment description

Supplementary Table SII | Variance decomposition

Supplementary Table SIII | QTL summary

Supplementary Data SI | Genotypes

Supplementary Data SII | Phenotypes

Supplementary Data SIII | Variance decomposition simulations

Supplementary Note SI | Supplementary Methods describing the partitioning of trait variation into its genetic components

**SUPPLEMENTARY FIGURE LEGENDS**

**Supplementary Figure S1 Growth phenotype distributions**. (**a**) Variation in degree of stress in the different environments for maximum growth rate (left panel) and total growth (right panel). Note that actual population doubling time (hours), and total growth (cells), are shown to allow a direct biological interpretation of values. The transformation, from normalized trait values to actual doubling times and total growth, was achieved by multiplying normalized values with the median control trait value and reversion of the log-transformation for each environment. (**b**) Frequency distribution of normalized phenotypes for POLs (blue) and estimated diploid F12 parents (gray).

**Supplementary Figure S2 Extent of genome-wide homozygosity has no impact on yeast growth**. Each point represents one POL in one environment (colors), with maximum growth rate (left panel) and total growth (right panel) in different panels. *x*-axis shows mean heterozygosity, across all the segregating sites in the genome. *y*-axis shows the normalized growth phenotype.

**Supplementary Figure S3 QTL maps for each trait**. Four QTL mapping approaches were used for each trait: nonadditive QTL mapping, using residuals from the estimated diploid F12 parents (top panel); dominance, testing for significance of the dominance term using linear mixed models (second panel from the top); additive QTL mapping, using estimated diploid F12 parents (third panel from the top); and finally, additive QTL mapping, using linear mixed models (bottom panel). *y*-axis = LOD score, *x*-axis = genetic position.

**Supplementary Figure S4 Phenotype distributions as a function of genotype for key QTLs**. (**a**) Left panel: distribution of total growth in allantoin, as a function of genotype composition at the QTL at chr. IX. Right panel: total growth in galactose, as a function of genotype composition at the QTL on chr. IV. (**b**) Tukey boxplots for growth traits as a function of genotype composition at the near universal chr. IX QTL, penetrating in all but one environment with antagonistic effects on maximum growth rate and total growth. Note: on average, homozygote WA is the strongest genotype for maximum growth rate. However, as shown (Fig S5) heterozygotes are heavily enriched among the best performing POLs. Thus, dominance, gives way to overdominance, in some genetic backgrounds.

**Supplementary Figure S5 Dominance, overdominance and underdominance in heterotic POLs**. (**a**) Frequency of POLs not significantly different from their corresponding estimated diploid parents (y-axis) as a function of different FDR *q*-values (x-axis) where the red label (0.01) indicates FDR *q*-value used for downstream analysis. (**b**) For each growth phenotype (black label = growth rate, red label = total growth) the genotype frequencies for best parent heterotic POLs (BPH), all POLs and worst parent heterotic POLs (WPH) at the pleiotropic chr. IX QTL. Best parent heterotic POLs for growth rate at this segregant site are significantly overrepresented for the heterozygous genotype compared to all POLs in most environments (*p* < 0.01, exception of NaCl, glycine and caffeine) and are significantly underrepresented for the NA homozygous genotype (exception of NaCl and isoleucine). Conversely, best parent heterotic POLs in total growth are in all environments overrepresented for homozygous NA (*χ*^2^ test, *p*<0.01) and underrepresented for the heterozygote. (**c**) The percentage of WPH explained by underdominance as a function of FDR.

